# Genetic Susceptibility to Hypertension Drives Brain Region-Specific Cellular Stress, Autophagic Failure, and Apoptosis in Sabra Rats

**DOI:** 10.64898/2026.11.21.768333

**Authors:** Yaseen Awad-Igbaria, Moti Ben-Shabat, Reem Sakas, Saher Abu-ata, Shifra Sela, Bella Gross, Yoram Yagil, Chana Yagil, Eilam Palzur

## Abstract

Hypertension is a significant risk factor for neurodegeneration. However, whether genetic vulnerability to high-blood pressure is associated with cellular dysfunction that exacerbates neurodegeneration in a specific brain region remains uncertain. Here, we evaluated changes in blood pressure following dietary salt-loading or a regular diet in Sabra-Normotensive (SBN/y) and Sabra-Hypertension-prone rats (SBH/y). We performed RT-PCR and immunofluorescence in the hippocampus (HPC) and basal ganglia (BG) tissues before salt-loading and after salt-loading (6 and 9 months) to examine the expression of proteins involved in stress response (Activating Transcription Factor 4-ATF4; Eukaryotic Initiation Factor 2-eIF2α), autophagy (Autophagy related 4A cysteine peptidase-Atg4a; light-chain3-LC3A/B; mammalian target of rapamycin complex 1-mTORC1; Beclin1) and apoptosis (Cas-3). The current results show that prior to the salt-diet, in the HPC and BG, there was overactivation of Cas-3 and a lower level of cleaved LC3A/B in SBH/y compared to SBN/y. However, in the HPC we observed a higher transcription of ATF4, and a lower transcription of mTORC1, Beclin1, and ATG4A (mRNA/Protein) in SBH/y. In contrast, in the BG, we observed an increase in eIF2α phosphorylation. Following salt-loading, SBH/y but not SBN/y developed hypertension. In hypertensive SBH/y, there was an increase in the activity of Cas-3, eIF2α phosphorylation, and downregulation of mTORC1 and Beclin1 transcription. Current results suggest that genetic susceptibility to hypertension is associated with stress response overactivation, and autophagy dysfunction, which might contribute to cellular damage accumulation, eventually triggering apoptosis. Critically, however, the cellular dysfunction in both brain regions (HPC and BG) was exacerbated by salt-diet that concurrently induces the development of hypertension.

## 1. Introduction

Hypertension (HTN) is one of the most critical risk factors for heart disease, stroke, other cardiovascular diseases (CVDs), dementia, and cognitive impairment [1,2]. Globally, high blood pressure affects 30% to 45% of the population [3], and the prevalence is rising globally owing to the aging of the population and increases in exposure to lifestyle risk factors including: smoking, high sodium consumption, overweight, and lack of physical activity [4–7].

Previous evidence suggests that the interaction between environmental risk factors and genetic predisposition to HTN is crucial in developing high blood pressure [8,9]. Consequently, the combined impact of genetic factors and environmental risk factors is more influential in promoting high blood pressure than the effects of either factor alone [10], emphasizing the importance of considering genetic and environmental factors synergism in understanding their crucial role in the development of HTN.

HTN is a complex systemic condition with implications beyond its association with heart disease [11]. It is recognized as a risk factor for developing neurodegenerative diseases that manifest in dementia [12,13]. Thus, a substantial body of evidence has shown that high blood pressure is associated with impaired neuronal function that promote neuronal death, and neurodegeneration [10,14–16]. Previously, we have demonstrated that high blood pressure in a rat model of hypertension can leads neuronal damage in the cortex, including disruptions of the autophagy process, imbalances in gene-expression regulation, an increase in gene-translation repression, and neuronal demyelination[10,14]. Critically however, these changes were associated a significant rise in cortical apoptosis and neuronal death in rats with HTN [10,14].

Interestingly, previous studies have shown that the hippocampus (HPC), which is involved in learning and memory, is susceptible to blood pressure changes [17,18] due to the high metabolic demands that are supported by a vast microvascular density [19]. Thus, a slight increase in blood pressure can cause a significant damage to the cell function and subsequent neuronal loss. Hance might explain the reduction in hippocampal volume, which manifests in impaired memory performance observed among individuals with high blood pressure [18,20]. Moreover, HTN can potentially induce damage to the basal ganglia (BG), a neural network that plays a crucial role in motor behaviors and procedural memory [21]. Several findings indicate that the BG is susceptible to stroke, microbleeds, and atherosclerosis due to HTN [22–24]. The BG sensitivity might be attributed to the small arteries that supply the BG, which are vulnerable to mechanical stresses associated with HTN [22]. Although the impact of HTN on the HPC and the BG brain regions has been extensively studied, it remains unclear whether the damage caused by HTN is comparable between these regions.

The development of HTN results from a complex interaction between genetic and environmental factors. It is still unclear, however, whether genetic susceptibility to high blood pressure may already be associated with neuronal cell dysfunction involved in neurodegeneration development in different brain regions (HPC, BG). In the current study, we investigated in an inbred rodent model genetically predisposed to salt-sensitive HTN the long-term effect of an environmental factor (high salt dietary intervention) on an array of neuronal cell functions that have been related to neurodegeneration in the hippocampus and basal ganglia.

## 2. Material Methods

### 2.1. Animals

Male Sabra Hypertension-prone (SBH/y; Three-month-old) and Sabra Hypertensive-resistant (SBN/y; Three-month-old) rats were used in the current study as a model of salt-sensitive or resistant hypertension, respectively [25,26]. All animal procedures were approved by the Bar-Ilan University Animal Care Committee and were carried out in accordance with the National Institutes of Health Guide for the Care and Use of Laboratory Animals. During the study, animals were housed in groups of 3-4 rats in a sterilized solid bottom cage with contact bedding under controlled temperature and a 12:12 h light/dark cycle.

### 2.2. Animal model of HTN

Animals were salt loaded by feeding standard rat chow enriched with 8% NaCl (Cat. TD.92012, Harlan, Israel). SBH/y fed a high salt diet invariably develops HTN [25,26]. Four groups of animals were studied: SBH/y^(HSD)^ provided a high salt diet (HSD) as the experimental hypertensive group, SBH/y^(RD)^ provided a regular rat diet (RD) as the hypertension-prone salt-control group, SBN/y^(HSD)^ provided HSD as the hypertension-resistant salt-control group and SBN/y^(RD)^ on regular diet as the strain control group (**Fig.1a**). In addition, rats were provided with RD prior to switching to HSD.

### 2.3. Blood pressure measurements

Blood pressure was measured using a non-invasive tail-cuff method (CODA™, Kent Scientific Corporation, Torrington, CT, USA). Briefly, rats were placed in restraint tubes and left for 20 min to warm (tail skin surface temperature ∼30°C) on the instrument warming platform, and a pressure transducer was placed on the rat’s tail. To prevent stress, rats were allowed to habituate to this procedure for 5 days before measurements were performed. The blood pressure was averaged from 10 consecutive readings. The Coda^TM^ instrument displays diastolic, systolic, and mean arterial blood pressure (MAP). However, we used the mean arterial blood pressure (MAP) for the analyses. The blood pressure test was performed by an observer blinded to the condition.

### 2.4. Histology and immunohistochemistry

Animals were deeply anesthetized, and transcardially perfused with heparinized saline, 10% sucrose in buffered saline, and 4% buffered formaldehyde[27]. The brain was excised and fixated in 5% paraformaldehyde in phosphate buffer for one hour and then embedded in paraffin. Sections of 5μm thickness were cut from blocks containing the hippocampus, or the basal ganglia using a microtome. The sections were then mounted on a Matsunami adhesive glass slide (TM-1190 TOMO, MATSUNAMI).

### 2.5. Immunohistochemistry

Immunofluorescent staining was performed as described before [28–30]. Briefly, slides were then deparaffinized and subjected to heat-induced epitope retrieval. To remove background staining, we used Background Buster (#NB306-50, INNOVEX), followed by three cycles of wash buffer, then incubated for one hour at room temperature in a humidity chamber with one of the antibodies: anti-ATG4A (1:200; #EPR4122, Millipore); anti-MAP1LC3A/B (1:200; #SAB4300, Novus Biologicals); cleaved anti-Caspase-3 (1:100; #CPP324, Zytomed System); anti-eIF2α∼P (1:200; TA343946 OriGene Inc.). After that, the slides were washed three times and incubated for 1 hour with a secondary antibody: anti-rabbit/mice 594 (1:150; Bethyl Laboratories, Inc.), followed by three cycles of wash buffer. Next, the sections were mounted with “Flouromount G with DAPI” (eBiosciences) and incubated for 30 seconds.

### 2.6. Microscopy

Microscopic analyses were performed using the Eclipse Ci microscope (Nikon Corp., Japan). Ten-images were captured by the Nikon DS-Ri1 camera (Nikon Corp., Japan) with the same microscope settings nd analyzed using the Image J software 1.46 by an observer blinded to the condition.

### 2.7. Gene expression analysis

Animals were sacrificed, the HPC and the BG were isolated and kept at –80° C. The RNA was extracted from the HPC and the BG by combining TRI reagent (Sigma-Aldrich) and Purelink TM kit (Thermo Fisher Scientific). cDNA was prepared, and real-time PCR was performed as described before[31–33]. The relative expression of the target genes normalized to glyceraldehyde-3-phosphate dehydrogenase (GAPDH) was calculated using the ΔCt method. The Gene primer sequences are listed in Table S1 (See Supplemental Material Table S1).

### 2.8. Data analysis

Statistical analyses were performed using IBM SPSS version 26 and GraphPad Prism. All data were expressed as Mean ± SEM. Differences between groups were assessed by Student *t*– test, one-way ANOVA, and three-way ANOVA. Differences were determined Post hoc by Tukey’s test and Student *t*-test when significant main effects or interactions were detected. The significance level was set at *P*<0.05.

## 3. Results

### 3.1. Blood pressure

Only the strain with genetic predisposition to HTN develop high blood pressure after oner month of HSD. Notable, the MAP of SBH/y^(HSD)^ was significantly higher at each time point compared to SBN/y^(RD/HSD)^ and SBH/y^(RD)^ (*P*<.00, **Fig. 1c**). There was no significant difference in MAP between SBN/y^(HSD)^ group and SBN/y^(RD)^ and SBH/y^(RD)^ treated with RD (See Supplemental Material Table S2).

**Figure. 1.**
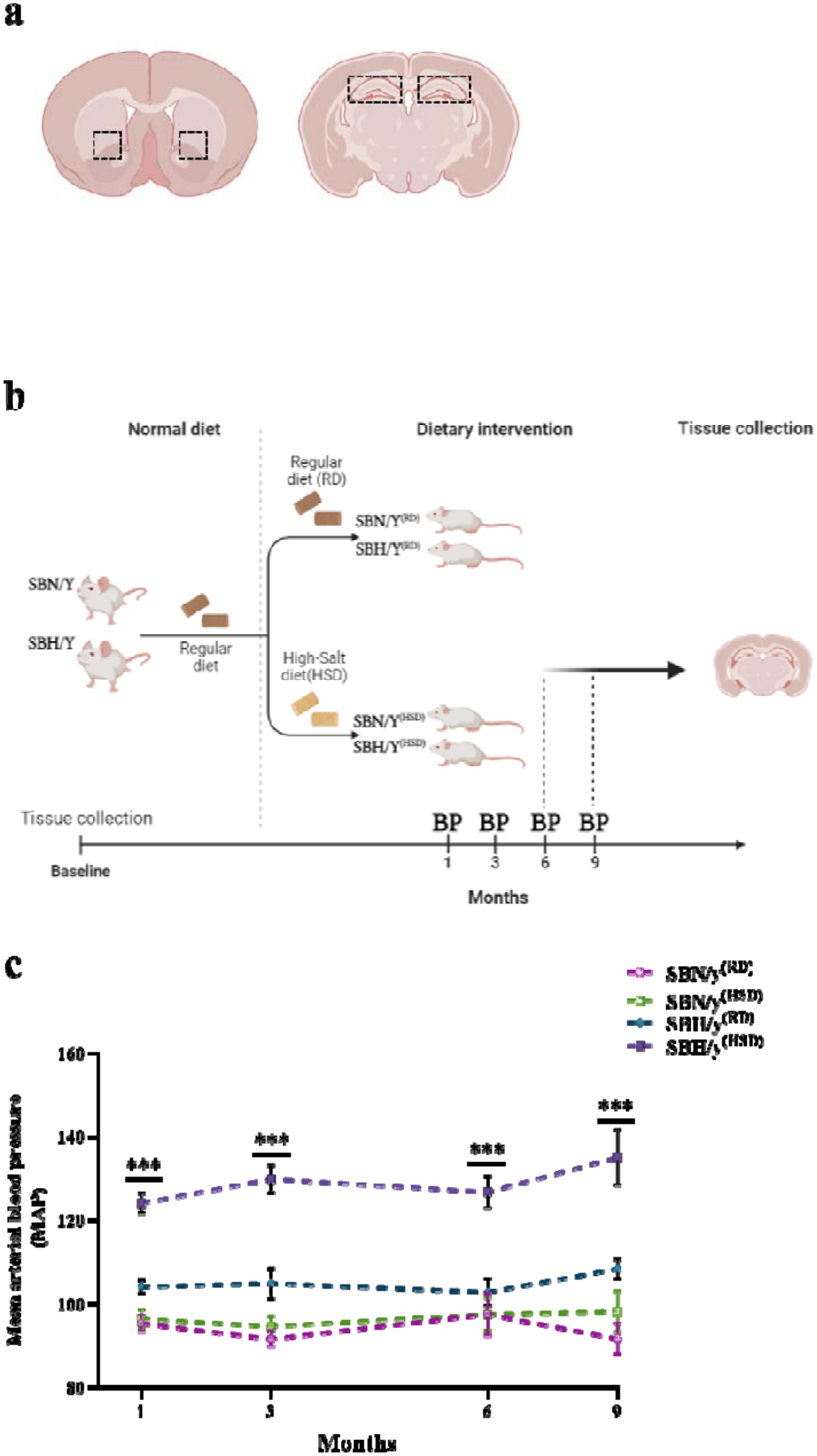
Schematic brain section and the experiment timeline. **(a)** Schematic coronal brain section of the hippocampus and the basal ganglia region where RTPCR and immunohistochemistry staining were done. **(b)** The timeline illustrates the experimental procedures across 9 months of the study; the Animals were separated into four groups: SBN/y^(RD)^ and SBH/y^(RD)^, provided a regular rat diet (RD), SBN/y^(HSD)^, and SBH/y^(HSD)^ provided a high salt diet (HSD). Blood pressure was measured at 4 points: 1, 3, 6, and 9 months after diet loading. Animals were sacrificed at three time points: baseline, 6 months, and 9 months after diet loading. **(c)** MAP of the four groups (n=4-17, per group). Mean ± SEM. One-way ANOVA-followed by Tukey’s multiple comparison test. \*\*\**P<*0.001, SBH/y+ vs. other groups.

### 3.2. Gene and protein expression profile of autophagy, stress response, and cell death

Prior to dietary intervention, we examined the differences in gene expression between SBH/y and SBN/y. In the HPC, the analysis revealed a significantly lower expression of Beclin-1, mTORC-1, Atg4A, and higher expression of ATF4 in SBH/y compared to SBN/y (**Fig. 2a**). While, in the BG, we only observed a significant increase in the expression of caspase-3 in SBH/y compared to SBN/y (**Fig. 2a**).

**Figure. 2.**
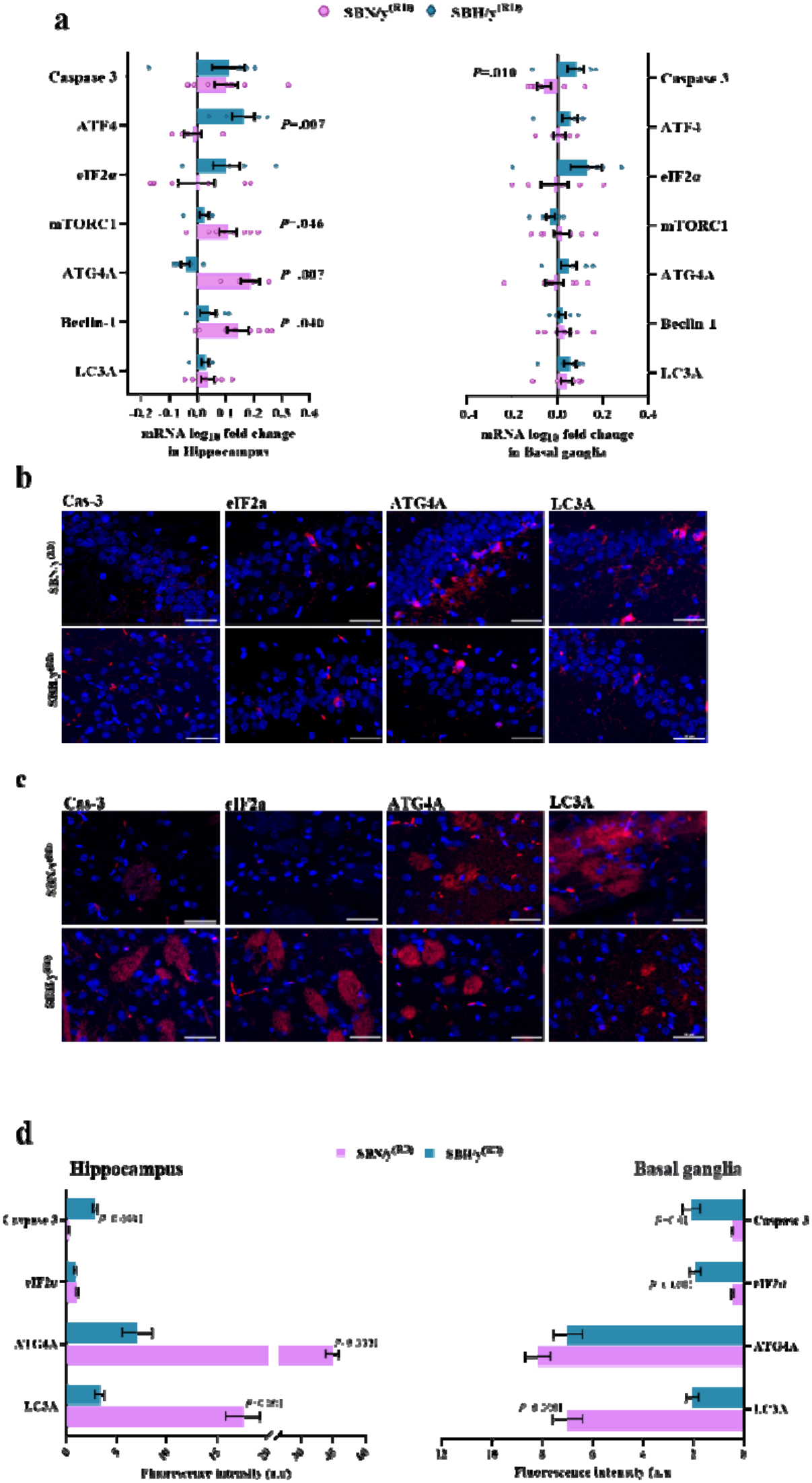
Gene and protein expression in the HPC and BG. **(a)** The expression of gene related to autophagy, stress response and apoptosis in the hippocampus and the basal ganglia (Fold change log 10, n=6-8 per group). **(b)** Representative merged images of DAPI – DNA blue staining of cells’ nuclei and protein related to autophagy (ATG4A and LC3A), stress response (eIF2a) and apoptosis (Caspace3) in hippocampus (red staining) in SBN/y and SBH/y at the baseline before diet loading. *Scale bar: 50 µm*. **(c)** Representative merged images of DAPI – DNA blue staining of cells’ nuclei and protein related autophagy (ATG4A and LC3A), stress response (eIF2a) and apoptosis (Caspace3) in the basal ganglia (red staining) in SBN/y and SBH/y at the baseline before diet loading. *Scale bar: 50 µm*. **(d)** IF analysis of the protein expression level in the HPC and BG (n=6-7 per group). Mean± SEM. Student t-test.

To obtain a more comprehensive understanding, we conducted a protein expression analysis in both brain areas: hippocampus and basal ganglia, to supplement our previous examination of mRNA expression differences between the SBH/y and SBN/y prior to dietary intervention. In HPC and BG, the analysis revealed a significant reduction in the expression of the cleaved form of microtubule-associated protein 1 light chain 3 (MAP1LC3)-LC3 in SBH/y compared to SBN/y (**Fig. 2b-d**). Additionally, in both brain regions, we found a significant increase in the expression of cleaved caspase 3 in SBH/y compared to SBN/y (**Fig. 2b-d**). However, reduced hippocampal Atg4A was noted in SBH/y compared to SBN/y, with no difference in the Atg4A expression in the BG between groups (**Fig. 2b-d**). On the other hand, a higher level of the phosphorylated eIF2α in SBH/y compared to the SBN/y was only observed in the BG but not in the HPC (**Fig. 2b-d**). The current result suggests that brain region might matter in the association between genetic predisposition to HTN and cellular dysfunction in critical cellular processes. However, whether the genetic predisposition to salt-induced hypertension might exacerbate the cellular dysfunction in both brain regions is unclear.

### 3.3. Autophagy process in the HPC and BG

We found that strain significantly impacted the expression of Atg4A at the mRNA level in the HPC. The SBH/y^(RD/HSD)^ groups had lower levels of mRNA-Atg4A compared to the SBN/y^(RD/HSD)^ groups at both 6 and 9 months after diet-loading (**Fig. 3a**). In the BG, there was also a significantly lower expression of mRNA-Atg4A in SBH/y^(HSD)^ compared to SBN/y^(HSD)^ after 9 months of diet-loading. Additionally, it was found that the expression of mRNA-Atg4A in the SBH/y^(HSD)^ group was significantly lower after 9 months of diet-loading in both brain regions (HPC and BG) compared to 6 months (*P<*0.001; **Fig. 3a**).

**Figure. 3.**
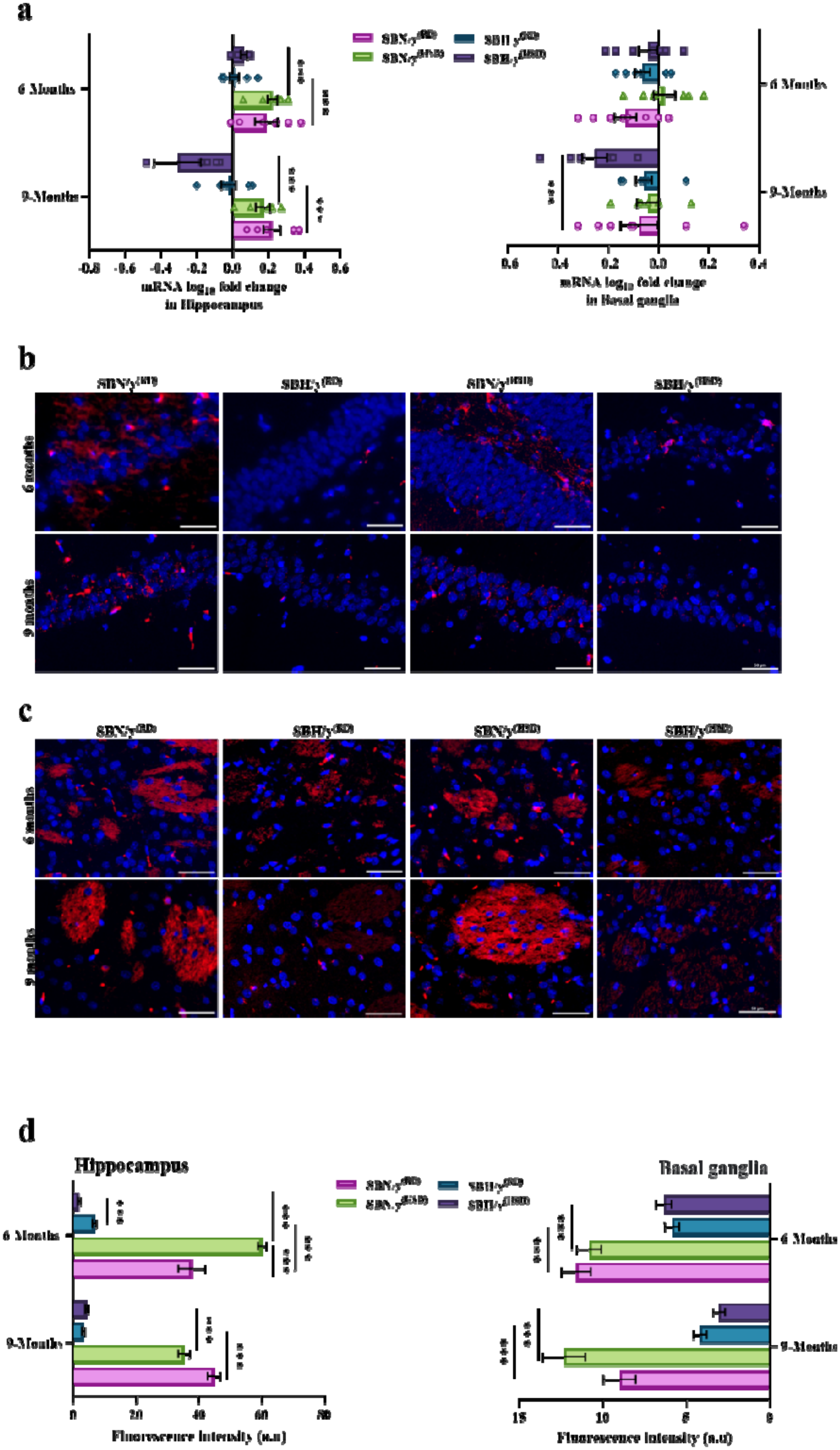
The expression of Cysteine protease (ATG4A) at the mRNA and protein level in the hippocampus and basal ganglia. **(a)** Cysteine protease (ATG4A) expression at the mRNA level in the hippocampus and basal ganglia of SBN/y and SBH/y at two-time points: 6 months, and 9 months after diet loading (n=6-8 per group). **(b)** Representative merged images of DAPI – DNA blue staining of cells’ nuclei and hippocampus-Atg4a protein-red staining in SBN/y and SBH/y at two-time points: 6 months, and 9 months after diet loading. *Scale bar: 50 µm*. **(c)** Representative merged images of DAPI – DNA blue staining of cells’ nuclei and basal ganglia-Atg4a protein-red staining in SBN/y and SBH/y at two-time points: 6 months, and 9 months after diet loading. *Scale bar: 50 µm*. **(d)** IF analysis of Atg4a protein level in hippocampus and basal ganglia (n=6-7 per group). Mean± SEM. Three-way ANOVA was followed by student t-test. \*\**P<*0.005; \*\*\**P<*0.001.

At the protein level, strain significantly affected the expression of Atg4A in the HPC and BG (**Fig. 3b-d**). At each time point, the expression of Atg4A in SBH/y^(RD/HSD)^ groups was significantly lower than in SBN/y^(RD/HSD)^ groups (**Fig. 3b-d**). In the HPC, there was also a significant effect of diet duration. This effect was noted in SBN/y^(HSD)^ and SBH/y^(RD)^ by reduced expression of Atg4A and an elevated level in SBH/y^(HSD)^ after 9 months of diet-loading compared to 6 months (**Fig. 3b-d**). Notable, after 6 months of diet loading, higher expression of Atg4A was noted in SBN/y^(HSD)^ compared to SBN/y^(RD)^, and SBH/y^(RD)^ compared to SBH/y^(HSD)^ (**Fig. 3b-d**). On the other hand, in the BG, the effect of diet duration was noted in SBH/y^(HSD)^ by reduced expression of ATG4A after 9 months of diet-loading compared to 6 months (**Fig. 3b-d**).

Atg4A converts microtubule-associated protein 1 light chain 3 (MAP1LC3)-LC3 in autophagy to the cleaved form LC3A/B. Here, we measured changes in the expression of LC3A at the mRNA level and the post-translation modification level (Cleaved form-LC3A/B). In the HPC, there was only a significant effect of strain on the expression of mRNA-LC3A, thus at each time point SBH/y^(RD/HSD)^ groups expressed lower levels of mRNA-LC3A compared to the SBN/y^(RD/HSD)^ groups (*P*<0.001; **Fig. 4a**). In addition, the expression of mRNA-LC3A increased in SBN/y^(HSD)^ and decreased in SBH/y^(HSD)^ compared to the regular diet after 6 months of diet-loading (*P*<0.001; **Fig. 4a**). In the BG, there was no significant effect of strain, diet type or diet duration on the expression of mRNA-LC3A (**Fig. 4a**). However, there was a significant interaction between strain and diet type, this interaction was derived by lower expression of mRNA-LC3A in SBH/y^(HSD)^ compared to SBN/y^(HSD)^ after 6 months of diet-loading, and higher expression in SBH/y^(RD)^ after 9 months of diet-loading compared to SBN/y^(RD)^, and SBH/y^(RD)^ after 6 months of diet-loading (**Fig. 4a**).

**Figure. 4.**
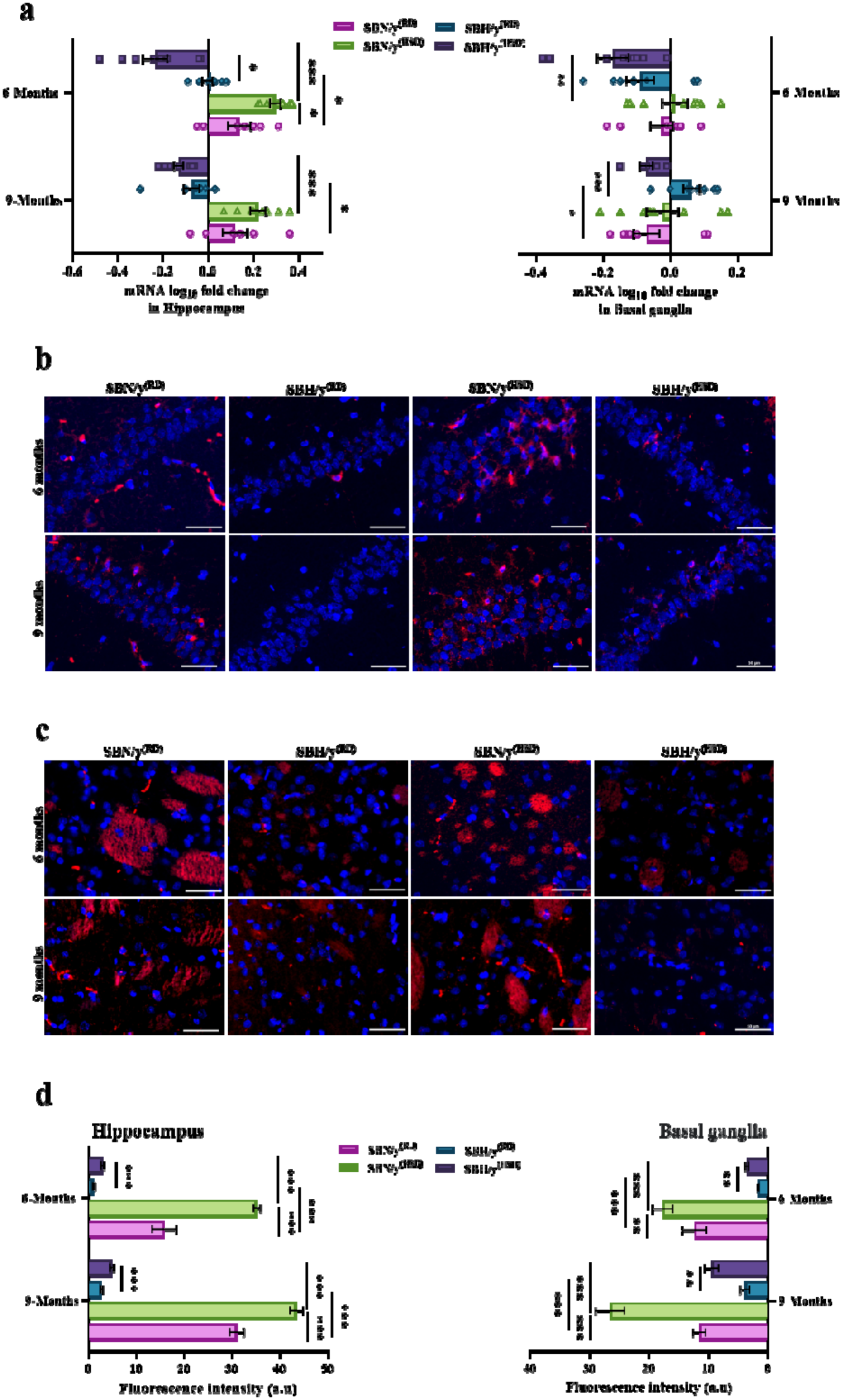
The expression of LC3A at the mRNA and protein level in the hippocampus and basal ganglia. **(a)** The expression of LC3A at the mRNA level in the hippocampus and basal ganglia of SBN/y and SBH/y at two-time points: 6 months, and 9 months after diet loading (n=6-8 per group). **(b)** Representative merged images of DAPI – DNA blue staining of cells’ nuclei and hippocampus-LC3A/B protein-red staining in SBN/y and SBH/y at two-time points: 6 months, and 9 months after diet loading. *Scale bar: 50 µm*. **(c)** Representative merged images of DAPI – DNA blue staining of cells’ nuclei and basal ganglia-LC3A/B protein-red staining in SBN/y and SBH/y at two-time points: 6 months, and 9 months after diet loading. *Scale bar: 50 µm*. **(d)** IF analysis of LC3A/B protein level in hippocampus and basal ganglia (n=6-7 per group). Mean± SEM. Three-way ANOVA was followed by followed by student t-test. \**P<*0.05; \*\**P<*0.005; \*\*\**P<*0.001.

At the post-translation modification level (Cleaved form), in the HPC and BG, there was a significant effect of strain, diet type, and diet duration on the expression of LC3A/B (*P*<0.001; **Fig. 4b-d**). At each time point, SBH/y provided HSD or RD expressed lower levels of LC3A/B protein than SBN/y^(RD/HSD)^ groups (*P*<0.001; **Fig. 4b-d**). In addition, HSD increased the expression of LC3A/B in both strains (SBH/y and SBN/y) compared to RD (*P*<0.001; **Fig. 4b-d**). The effect of diet duration was driven by increased expression of LC3A/B in all groups: SBH/y^(RD/HSD)^, SBN/y^(RD/HSD)^, after 9 months of diet-loading compared to 6 months (Excluding in the BG of SBN/y^(RD)^, and SBN/y^(HSD)^; *P*<0.001; **Fig. 4b-d**).

To further explore autophagy process in our model of salt-sensitive hypertension, we examined the expression of the regulators of autophagy – mTORC1 and Beclin1 at the mRNA level. In the HPC, there was a significant interaction and effect of strain, diet duration, on the expression of mTORC1 (**Fig. 5a**). Thus, at each time point after diet-loading (6 and 9 months) SBH/y^(RD/HSD)^ groups expressed lower level of mTORC1 compared to SBN/y^(RD/HSD)^ groups (*P*<0.05; **Fig. 5a**). The effect of diet duration was noted in SBH/y^(RD)^ and SBN/y^(HSD)^ by elevated expression of mTORC1, after 9 months of diet-loading compared to 6 months (**Fig. 5a**). The interaction was derived by lower expression of mTORC1 in SBH/y^(HSD)^ compared to SBH/y^(RD)^, and higher expression in SBN/y^(HSD)^ compared to SBN/y^(RD)^ after 9 months of diet loading (**Fig. 5a**).

**Figure. 5.**
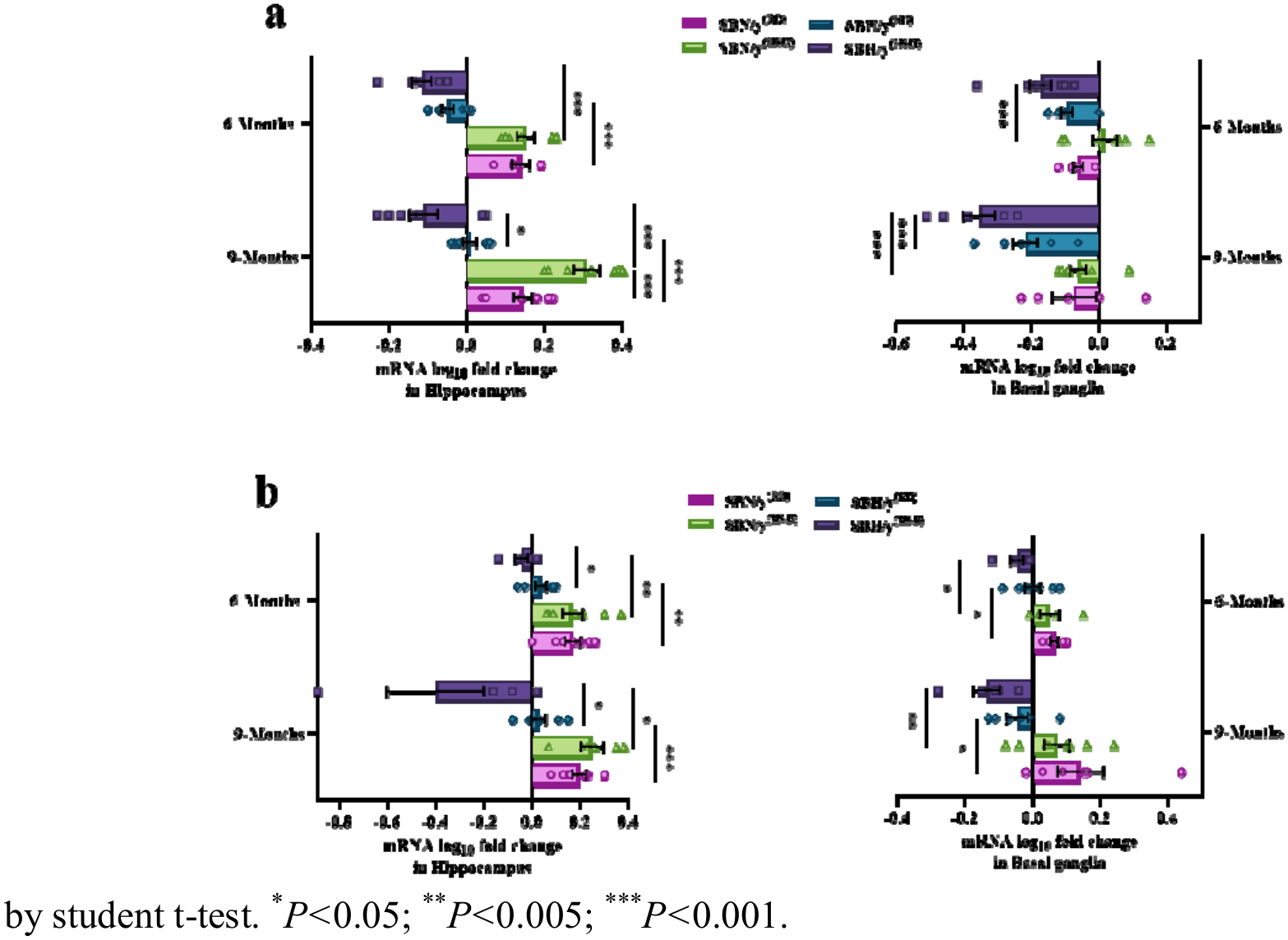
The expression of protein related to autophagy regulation process. **(a)** The expression of mTORC at the mRNA level in the hippocampus and basal ganglia of SBN/y and SBH/y at two-time points: 6 months, and 9 months after diet loading (n=6-8 per group). **(b)** The expression of Beclin-1 at the mRNA level in the hippocampus and basal ganglia of SBN/y and SBH/y at two-time points: 6 months, and 9 months after diet loading (n=6-8 per group). Mean± SEM. Three-way ANOVA was followed by student t-test. \**P<*0.05; \*\**P<*0.005; \*\*\**P<*0.001.

In the BG, there was a significant effect of strain, diet duration, on the expression of mTORC1 (**Fig. 5a**). Thus, at each time point after diet-loading (6 and 9 months) SBH/y^(HSD)^ expressed lower level of mTORC1 compared to SBN/y^(HSD)^ (*P*<0.05; **Fig. 5a**). The effect of diet duration was noted in SBH/y^(HSD)^ by reduction in the expression of mTORC1, after 9 months of diet-loading compared to 6 months (**Fig. 5a**). In addition, we noted interaction between strain and diet type, this interaction was derived by lower expression of mTORC1 in SBH/y^(HSD)^ compared to SBH/y^(RD)^ after 9 months of salt-diet loading (**Fig. 5a**).

Furthermore, in the HPC and BG there was significant effect of strain on the expression of Beclin1 at the mRNA level, thus at each time point after diet-loading (6 and 9 months), SBH/y^(RD/HSD)^ groups expressed lower level of mRNA-Beclin1 compared to SBN/y^(RD/HSD)^ groups (*P*<0.05; **Fig. 5b**). Nevertheless, the effect of diet type was noted only in the HPC, thus SBH/y^(HSD)^ expressed lower level of Beclin1 compared to SBH/y^(RD)^ at each time point after diet-loading (6 and 9 months) (**Fig. 5b**).

### 3.4. The effect of genetic predisposition to salt-sensitive hypertension on the stress response in the HPC and BG

In the BG, there was no significant effect of strain, diet type, and diet duration on mRNA-ATF4 expression (**Fig. 6a**). However, there was a significant effect of strain on mRNA-eIF2α expression (**Fig. 6a**), this effect was noted in SBH/y^(RD/HSD)^ groups by elevated expression of mRNA-eIF2α compared to SBN/y^(RD/HSD)^ after 6 months of diet-loading (**Fig. 6a**). Notable, a significant reduction of mRNA-eIF2α was noted in SBH/y^(HSD)^ after 9 months of diet-loading compared to 6 months (**Fig. 6a**).

**Figure. 6.**
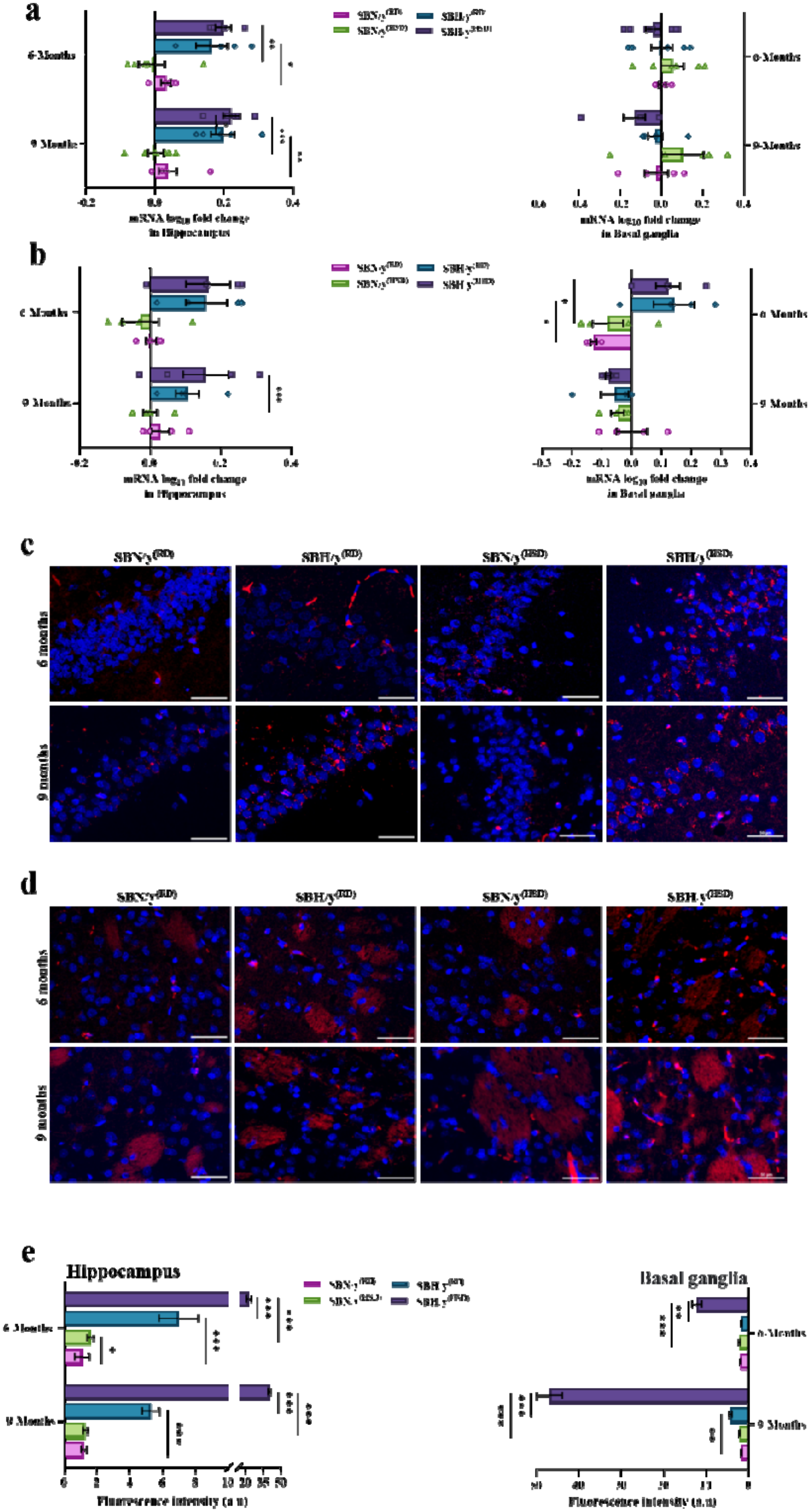
The expression of ATF4 and eukaryotic translation initiation factor 2 Alpha (eIF2α) at the mRNA level and phosphorylated (eIF2α∼P) at the protein level. **(a)** ATF4-mRNA expression in the hippocampus and basal ganglia of SBN/y and SBH/y at two-time points: 6 months, and 9 months on RD or HSD (N=6-8 per group). **(b)** eIF2α-mRNA expression in the hippocampus and basal ganglia of SBN/y and SBH/y at two-time points: 6 months, and 9 months after diet loading (n=6-8 per group). **(c)** Representative merged images of DAPI – DNA blue staining of cells’ nuclei and hippocampus-eIF2α∼P protein-red staining in SBN/y and SBH/y at two-time points: 6 months, and 9 after diet loading. *Scale bar: 50 µm*. **(d)** Representative merged images of DAPI – DNA blue staining of cells’ nuclei and basal ganglia-eIF2α∼P protein-red staining in SBN/y and SBH/y at two-time points: 6 months, and 9 months after diet loading. *Scale bar: 50 µm*. **(e)** IF analysis of eIF2α∼P protein level in hippocampus and basal ganglia (n=6-7 per group). Mean± SEM. Three-way ANOVA was followed by student t-test. \**P<*0.05; \*\**P<*0.005; \*\*\**P<*0.001.

On the other hand, in the HPC, there was only a significant effect of strain on the expression of ATF4 and eIF2α at the mRNA level (**Fig. 6a, b**). Thus, at each time point, there was a higher expression of mRNA-ATF4 in SBH/y^(RD/HSD)^ groups compared to SBN/y^(RD/HSD)^ groups (*P*<0.01; **Fig. 6a**), and a higher expression of mRNA-eIF2α in SBH/y^(HSD)^ compared to SBN/y^(HSD)^ after 9 months of diet-loading (**Fig. 6b**)

Since salt-loading affects protein activation by post-translational modifications, we examined whether eIF2α phosphorylation is altered during HSD. Subsequently, in both brain regions (HPC and BG) there was a significant interaction of strain, diet type, and diet duration on the level of eIF2α phosphorylation (**Fig. 6c-e**). In the HPC and BG, at each time point after diet-loading, SBH/y^(HSD)^ expressed higher levels of eIF2α phosphorylation than SBH/y^(RD)^ or SBN/y^(HSD)^ group (*P*<0.05; **Fig. 6c-e**). There was different pattern of change as a result of HSD in each brain region, thus in the HPC, at each time point the SBH/y^(RD/HSD)^ groups expressed higher levels of eIF2α phosphorylation compared to SBN/y^(RD/HSD)^ groups (*P*<0.05; **Fig. 6c-e**). Moreover, in the HPC, the interaction and the effect of diet duration was noted in SBH/y^(HSD)^ by elevated expression of eIF2α phosphorylation after 9 months of diet-loading compared to 6 months (*P*<0.05; **Fig. 6c-e**). On the other hand, in the BG, at each time point SBH/y^(RD/HSD)^ groups expressed higher levels of eIF2α phosphorylation compared to SBN/y^(RD/HSD)^ (Excluding 6 months after diet-loading between the RD groups; *P*<0.05; **Fig. 6c-e**). The interaction and the effect of diet duration was noted in SBH/y^(RD/HSD)^ by elevated expression of eIF2α phosphorylation after 9 months of diet-loading compared to 6 months (*P*<0.05; **Fig. 6c-e**).

### 3.5. The effect of genetic predisposition to salt-sensitive hypertension on apoptosis process

In the HPC there was no significant effect of strain, diet type, and diet duration on the expression of caspase-3 at the mRNA level (*P*<0.001; **Fig. 7a**). In the BG, there was only a significant effect of strain, which was driven by elevated expression of caspase-3 in SBH/y^(RD)^ compared to SBN/y^(RD)^ at each time point after diet-loading (**Fig. 7a**). At the protein level, there was a significant effect of strain, diet type, and diet duration on the expression of caspase-3 in the HPC and BG (**Fig. 7b-d**). At each time point, SBH/y^(RD/HSD)^ groups expressed higher levels of caspase-3 compared to the SBN/y^(RD/HSD)^ groups (*P*<0.001; **Fig. 7b-d**). Additionally, in the HPC and BG, HSD increased the expression of caspase-3 in SBH/y (*P*<0.001; **Fig. 7b-d**). However, only in the BG, HSD also increased the caspase-3 expression in SBN/y after 9 months of diet-loading (**Fig. 7b-d**). Finally, in the HPC and BG, caspase-3 expression was elevated after 9 months compared to 6 months only in SBH/y^(HSD)^ (*P*<0.001; **Fig. 7b-d**).

**Figure. 7.**
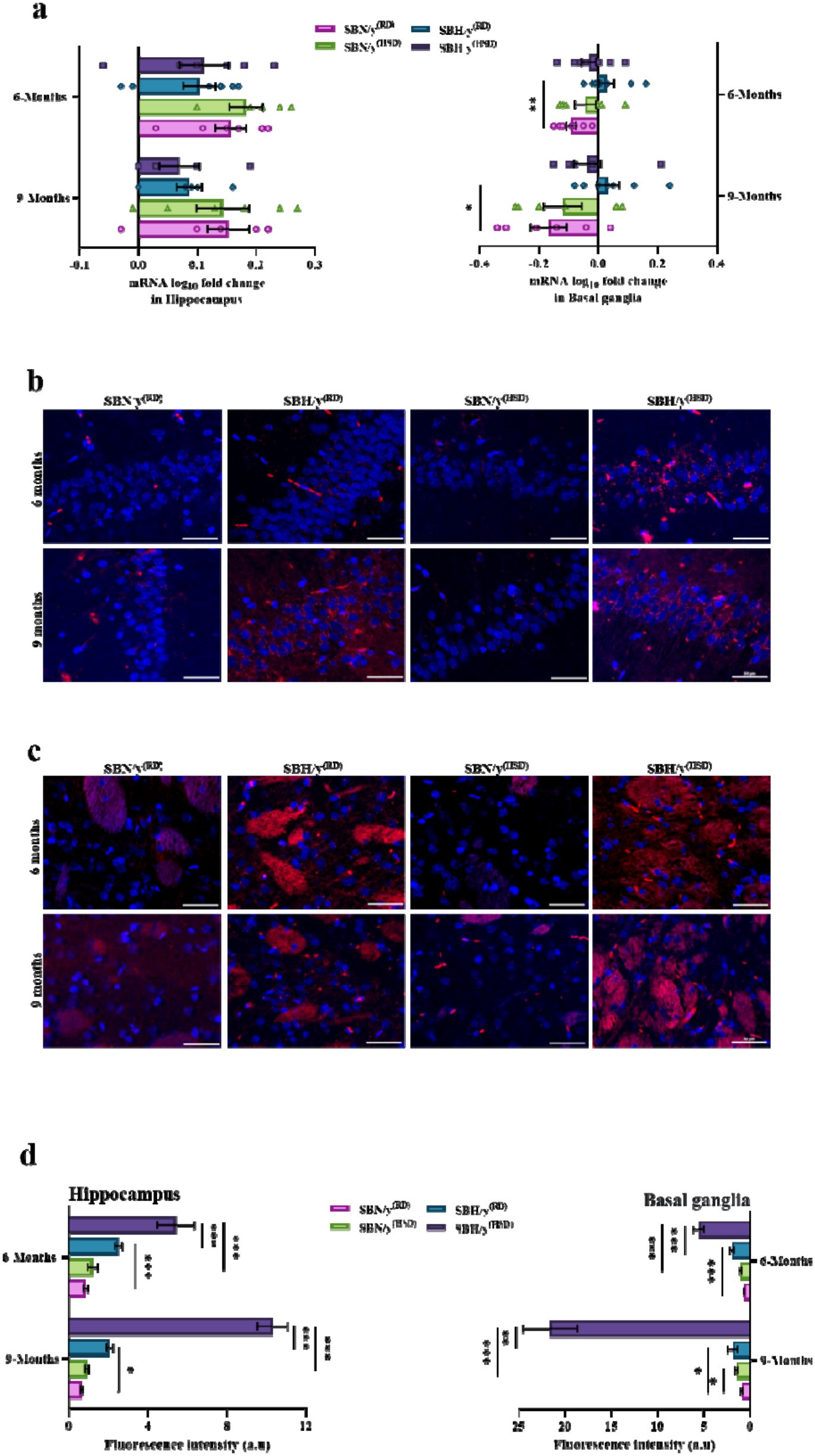
The expression of Caspase-3 at the mRNA and protein level in the hippocampus and basal ganglia. **(a)** The expression of caspase-3 at the mRNA level in the hippocampus and basal ganglia of SBN/y and SBH/y at two-time points: 6 months, and 9 months after diet loading (n=6-8 per group). **(b)** Representative merged images of DAPI – DNA blue staining of cells’ nuclei and hippocampus-caspase-3 protein-red staining in SBN/y and SBH/y at two-time points: 6 months, and 9 months after diet loading. *Scale bar: 50 µm*. **(c)** Representative merged images of DAPI – DNA blue staining of cells’ nuclei and basal ganglia-caspase-3 protein-red staining in SBN/y and SBH/y at two-time points: 6 months, and 9 months after diet loading. *Scale bar: 50 µm*. **(d)** IF analysis of caspase-3 protein level in hippocampus and basal ganglia (n=6-7 per group). Mean± SEM. Three-way ANOVA was followed by student t-test. \**P<*0.05; \*\**P<*0.005; \*\*\**P<*0.001.

**Figure 8.**
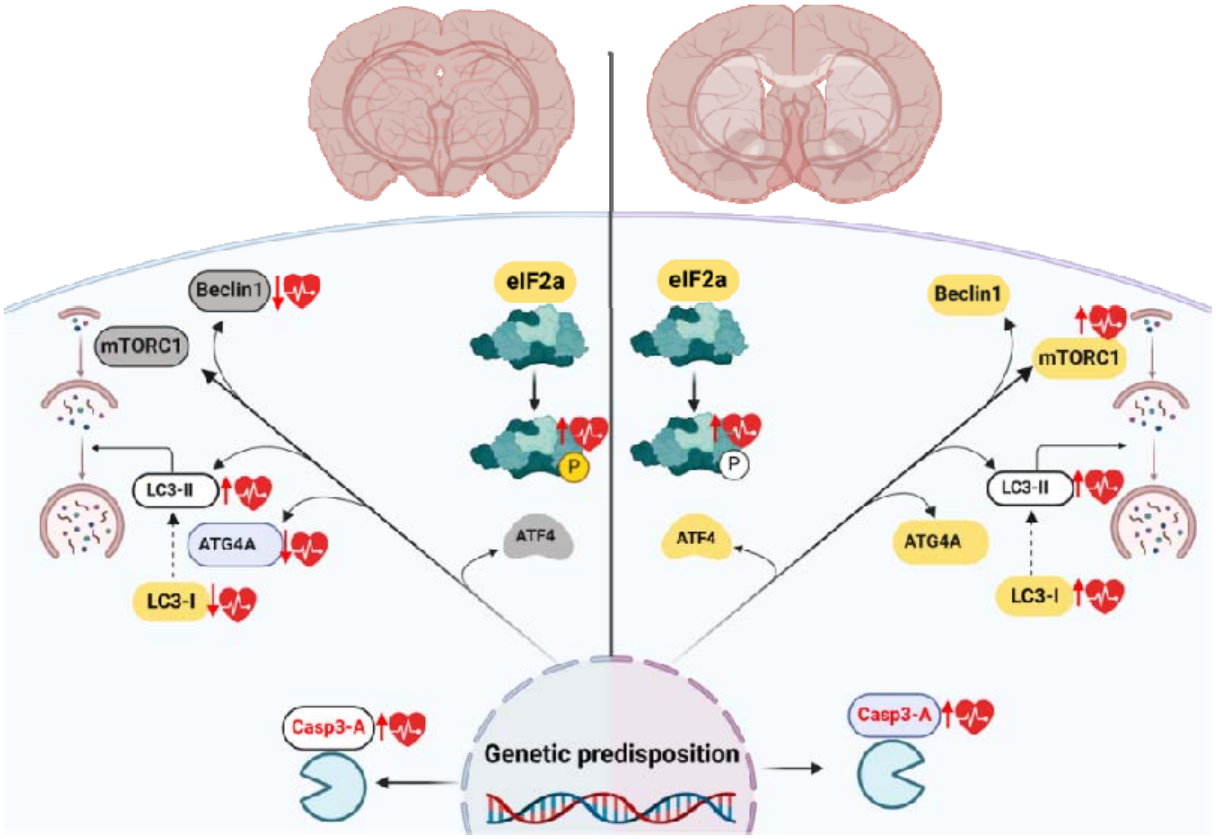
Diagram illustrating the molecular neurodegeneration machinery in hypertension in the SBH/y strain following salt loading. A genetic predisposition to salt-sensitive hypertension is linked to cellular dysfunctions in gene transcription regulation, autophagy, stress response, and apoptotic processes in the hippocampus and basal ganglia. These dysfunctions are further exacerbated by salt loading, which simultaneously induces hypertension. mRNA is represented in gray, proteins in white, and their combination in gold, while the effects of hypertension on gene expression or/and protein levels are indicated by a red heart.

## 4. Discussion

The current study shed light on several cellular factors promoting neurodegeneration and provides evidence for a neurodegenerative process secondary to HTN in a genetic rat model. The SBH/y strain has a genetic predisposition to develop high blood pressure when exposed to an HSD[26] with no evidence of spontaneous HTN development [26]. Our findings are consistent with previous reports[34]. Thus, we found that only SBH/y treated with a salt diet developed hypertension, whereas SBH/y provided RD, and SBN/y, irrespective of the diet, remained normotensive. More critically, here we found that the genetic predisposition of the SBH/y was already associated with a dysfunction in the autophagy process, cellular stress response, and elevated level of cell death in the HPC and BG.

Interestingly, our results indicate that the brain region affects the association between genetic susceptibility to HTN and dysfunction of the autophagy process and cellular stress response. In the HPC and BG, we found a significant reduction in converting microtubule-associated protein 1 light chain 3 (MAP1LC3)-LC3 to the cleaved form-LC3A/B in SBH/y compared to the SBN/y strain, despite comparable transcription levels between strains. However, the dysfunction of the autophagy process was more meaningful and widespread in the HPC compared to the BG in SBH/y. Specifically, we observed a reduction in the mRNA and protein levels of Atg4a and a decrease in the mRNA expression of autophagy regulators mTORC1 and Beclin1 only in the HPC region. These findings suggest that the HPC may be particularly vulnerable to the adverse effects of genetic susceptibility to HTN on the autophagy process.

In contrast, our results indicate that the BG region may be more susceptible to cellular stress than the HPC. Thus, we found a significant increase in the phosphorylated level of the translation repressor-eIF2α in the BG region, indicating that the cells in this region are under cellular stress compared to the HPC region. However, the increased transcription level of ATF4 in the HPC might be a sign of damage to the cellular response to stress. Together, these findings suggest that the relationship between genetic susceptibility to HTN and neuronal dysfunction is complex and varies across different brain regions.

The initiation factor eIF2α and the transcription factor ATF4 play a vital role in the cellular stress survival mechanism [35]. In response to the cellular stress, eIF2α phosphorylation represses global translation coincident with the preferential translation of ATF4 [36,37], which helps to alleviate the metabolic, oxidative, and ER stress by conserving nutrients, and promoting the expression of adaptive genes [38]. Nevertheless, eIF2α and ATF4 have a dual role, thus overactivation of this mechanism may induce apoptosis [39]. In the current study, we found that hypertension salt-induced among rats with a genetic susceptibility to HTN did not affect the transcription level of eIF2α and ATF4, thus there was no further expression in SBH/y^(HSD)^ compared to SBH/y^(RD)^. However, at each time point after diet-loading, we noted a higher expression of the hippocampal-ATF4 and eIF2α in the BG after 6 months of diet-loading in SBH/y groups compared to SBN/y groups. More importantly, at the protein level, we found a robust increase in the phosphorylation of eIF2α in both brain regions (HPC and BG) among hypertensive rats treated with salt diet.

It should be noted that after six months, there was a slight increase in the phosphorylation of eIF2α in the HPC of the SBN/y group treated with a salt diet. However, the expression of eIF2α phosphorylated was higher in the SBH/y^(HSD)^ group than in the SBN/y^(HSD)^. Current results are consistent with the notion that HTN causes chronic neuronal stress, leading to excessive phosphorylation of eukaryotic translation initiation factor 2α (eIF2α-P), which eventually might impair neuronal function and contribute to neurodegeneration [40–42]. Furthermore, the current result underscores that the combined effects of an environmental factor (Salt-diet) and genetic susceptibility to HTN are more significant than the effects of each factor separately in provoking neuronal stress.

Previous studies have shown that HTN is associated with a robust increase in autophagy in neuronal cells [41,43]. The suggested mechanism hypothesizes that autophagy acts as a compensatory and adaptive response to stress by providing a dynamic homeostasis of nutrients in cells by degrading and recycling misfolded proteins [44]. Nevertheless, uncontrolled over-activation of autophagy could lead to the depletion of essential proteins required for the survival of cells, ultimately causing an increase in autophagic cell death [45]. Unexpectedly, we found that the genetic predisposition to salt-induced HTN was associated with a broad dysfunction in the autophagy process. First, lower levels of Atg4a protease were found in the HPC and BG of SBH/y compared to SBN/y after 6 and 9 months of diet-loading in both diet-type.

Furthermore, with aging, the expression of Atg4a was even more diminished in SBH/y, particularly in the HPC. The salt-loading has a minor effect on the expression pattern in both strains, thus after 6 months of salt-loading, we noted an increase in the expression of the hippocampal Atg4a in the SBN/y, and a slight decrease in the SBH/y provided with salt-diet compared to the regular diet groups, however, the differences was less detectable after 9 months of diet-loading. It’s possible that the observed changes in the expression of Atg4a may be attributed to a cellular adaptation process in response to a high-salt diet that occurs gradually over time, and probably this adaptation process is influenced by aging as well.

The diminished expression of Atg4a might reflect in reducing the autophagy cascades since Atg4a initiates the autophagy process by converting microtubule-associated protein 1 light chain 3 (MAP1LC3)-LC3 to the cleaved form-LC3A/B [46]. Despite comparable transcription levels of LC3A between strains at the baseline, after 6 and 9 months we found that SBH/y provided RD or HSD expressed lower level of mRNA-LC3A in the HPC. Yet, in the BG, only the SBH/y provided HSD shows reduction of mRNA-LC3A after 6 months of diet-loading. Notably, the transcription level was affected by the salt diet, thus in the HPC there was further transcription of LC3A in SBN/y^(HSD)^ and diminished in SBH/y^(HSD)^ after 6 months of diet-loading. Since the transcriptional landscape does not necessarily reflect the protein levels, we measured the expression of the cleaved form-LC3A/B. Strikingly, in the HPC and BG, we found a diminished expression of the cleaved protein form LC3A/B in SBH/y at each time point. Additionally, in both strains, SBH/y and SBN/y, salt loading increased the level of the cleaved form LC3A/B. The putative explanation for the upregulation of autophagy may be related to the activation of corrective mechanisms such as autophagy, which aim to prevent accumulating cellular damage in neurons caused by salt [47,48]. On the other hand, when the autophagy process is defective, as in SBH/y, salt-diet induced-hypertension could cause chronic cellular stress in neurons that contribute to cell death.

Furthermore, our result indicates a decrease in the regulation of the autophagy process in the strain with a genetic predisposition to hypertension. In the HPC, the diminished expression of the autophagy regulators, mTORC1 and Beclin1, was directly associated with the genetic predisposition to hypertension, and salt-loading further reduce the expression. Nevertheless, in the BG, the expression of mTORC1 and Beclin1 was not directly associated with the genetic predisposition to hypertension and was mainly affected by the interaction between salt-diet and aging in SBH/y. Thus, at the baseline, no difference was noted between SBH/y and SBN/y. However, after 6 and 9 months of diet-loading a reduction in Beclin1 expression was noted in SBH/y groups with no further effect of salt-loading, in contrast, the reduced expression of mTORC1 was mainly ascribed to the salt-diet (SBH/y^(HSD)^). However, long-term consumption of a high-salt diet led to an increased expression of mTORC1 in SBN/y, this might be associated with the fact that salt-diet upregulate the autophagy process as a compensatory mechanism, which in turn requires more regulation to maintain homeostasis. Nevertheless, the reduction in mTORC1 and Beclin1 in hypertension rats may suggest that autophagy regulation is disrupted, independent of the brain region [49].

The overactivation of the gene regulation/expression mechanisms under stress, the dysfunction and the lack of autophagy regulation might contribute to the accumulation of cellular damage in neurons, which in turn trigger apoptosis. At the transcription level, an elevated level of caspase-3 was noted only in the BG of SBH/y strain. Critically however, in both brain regions (HPC and BG), we found a significant increase in the activated form of caspase-3 that was further amplified by salt-loading in rats with a genetic predisposition. It should be noted that a prolonged duration of a high-salt diet may cause cellular stress, which could result in a slight activation of caspase-3 in normotensive rats. Taken together, current results indicate that genetic predisposition to HTN is associated with elevated levels of apoptosis, which are further exacerbated by salt-induced hypertension. These findings are compatible with previous reports showing that HTN increases neuronal apoptosis, resulting in neurodegeneration [50].

In conclusion, our findings indicate that genetic susceptibility to HTN is associated with the overactivation of the cellular stress survival mechanism, dysfunction and the lack of autophagy regulation, which might contribute to the accumulation of cellular damage in neurons, eventually triggering apoptosis. Additionally, the brain region was found to affect this association, thus the dysfunction of the autophagy process was more meaningful and widespread in the HPC compared to the BG. In contrast, the BG region was more susceptible to activating the cellular stress response. Critically however, the cellular dysfunction in both brain regions (HPC and BG) was exacerbated by salt loading that concurrently induces the development of high blood pressure. Consequently, the genetic predisposition to cellular dysfunction thus appears to fulfill a significant primary role in the accumulation of cellular damage in neurons, whereas hypertension probably plays a secondary role by accelerating the process, which is already prevalent in the genetically susceptible.

## Abbreviations

HTN, Hypertension; HPC, hippocampus; BG, Basal Ganglia; ATF4, Activating Transcription Factor 4-ATF4; eIF2α, Eukaryotic Initiation Factor 2; autophagy (Autophagy related 4A cysteine peptidase-Atg4a; LC3A/B, light-chain 3; mTORC1, mammalian target of rapamycin complex 1; SBH/y, Sabra Hypertension-prone; SBN/y, Sabra Hypertensive-resistant; HSD, High salt diet; RD, Regular diet.

## Declaration

### Funding

The authors declare that no funds, grants, or other support were received during the preparation of this manuscript.

### Conflicts of Interest

The authors have no relevant financial or non-financial interests to disclose.

### Author contributions

Conceptualization, M.B.S., and Y.A.I..; Methodology, Y.Y., C.Y., S.S., Y.Y., S.A.A., and B.G.; Investigation, Y.A.I., M.B.S., R.S., S.A.A., Y.Y., and E.P.; Formal Analysis, Y.A.I.,M.B.S., and R.S.,; Resources, Y.Y., C.Y., S.S., B.G., and E.P.; Writing – Original Draft, Y.A.I., and M.B.S.; Writing – Review & Editing, S.S., and E.P.; Supervision, E.P.; Project Administration, Y.Y., C.Y., S.S., B.G., and E.P.

### Availability of data and materials

The datasets analyzed during the current study are available from the corresponding author upon request.

### Ethics approval

All animal procedures were approved by the Bar-Ilan University Animal Care Committee and were carried out in accordance with the National Institutes of Health Guide for the Care and Use of Laboratory Animals.

### Funding

The authors declare that no funds, grants, or other support were received during the preparation of this manuscript.

## Acknowledgements

The authors would like to thank Prof. Jean Francois Soustiel, Associate Dean at the Azrieli Faculty of Medicine for his support in research logistics and equipment.

## Notes

### Competing Interest Statement

The authors have declared no competing interest.

