## Supplemental Material Table S2 for "Genetic Susceptibility to Hypertension Drives Brain Region-Specific Cellular Stress, Autophagic Failure, and Apoptosis in Sabra Rats"

**Table S2. Mean Arterial Blood Pressure**

|  | SBN/y^(RD)^ | SBN/y^(HSD)^ | SBH/y^(RD)^ | SBH/y^(HSD)^ |
| --- | --- | --- | --- | --- |
| Months |  |  |  |  |
| 1 | 95.3±1.9 | 96.5±2.3 | 103.3±1.3 | 124.2±2.4^***^ |
| 2 | 99.8±3.2 | 97.9±2.0 | 103.9±1.0 | 124.4±1.4^***^ |
| 3 | 91.7±1.8 | 94.7±2.4 | 103.3±3.8 | 129.9±3.3^***^ |
| 4 | 96.8±2.8 | 97.6±2.3 | 106.9±1.9 | 126.8±2.5^***^ |
| 5 | 96.4±3.3 | 100.3±3.5 | 113.5±5.0 | 136.4±4.8^***^ |
| 6 | 97.6±5.1 | 97.6±4.0 | 102.9±3.3 | 126.8±3.8^***^ |
| 7 | 94.0±3.1 | 92.7±3.7 | 105.8±5.7 | 133.4±4.2^***^ |
| 8 | 94.6±4.3 | 94.8±4.4 | 105.6±3.1 | 130.1±4.2^***^ |
| 9 | 91.8±3.6 | 98.2±4.9 | 108.5±2.4 | 135.2±6.7^***^ |
| Mean±SEM | | | | |
