## Supplemental Material Table S1 for "Genetic Susceptibility to Hypertension Drives Brain Region-Specific Cellular Stress, Autophagic Failure, and Apoptosis in Sabra Rats"

**Table S1.** **The Gene primer sequences.**

| Primer name | F/R | Sequenc 5' - 3' | Accession of mRNA: RefSeq |
| --- | --- | --- | --- |
| Atg4a | F | ACAGTGCCTATAAAATAGGTGACGAG | NM_001126298 |
|  | R | AATCCCATAAAAGCTCCTTGACC |  |
| MAP1LC3A | F | AAGATCCCGGTGATCATCGA | NM_199500.2 |
|  | R | GCGCCGGATGATCTTGAC |  |
| mTORC1 | F | CCTCAGTGGTACAGGCACACA | NM_019906 |
|  | R | CTCAGACGCTCTCCCTCCAT |  |
| Becn1 | F | TGGACCAGGAGGAAGCTCAG | NM_001034117 |
|  | R | TGTCCGCTGTGCCAGATATG |  |
| Caspase-3 | F | CTCTTCATCATTCAGGCCTGC | NM_012922.2 |
|  | R | AGTAACCGGGTGCGGTAGAGTA |  |
| ATF4 | F | CATGGCGCTCTTCACGAAA | NM_024403.2 |
|  | R | GCTGGTATCGAGGAATGTGCTT |  |
| eIF2α | F | GTTCCAAGCTGCATGAAGAGG | NM_001109339.1 |
|  | R | TTCTTGCCTCCTGTTTTGCAG |  |
| GAPDH | F | AGAGACAGCCGCATCTTCTTG | NM_017008.4 |
|  | R | GTAACCAGGCGTCCGATACG |  |
